# De Novo Genome Assembly and Annotation for the Webbing Clothes Moth (*Tineola bisselliella*): A Globally Distributed, Economically Important Pest

**DOI:** 10.1101/2024.08.19.604795

**Authors:** Jasmine D Alqassar, Hannah E Aichelman, Isabel A Novick, Sean P Mullen

**Author notes:** Co- corresponding authors (;).

## Abstract

*Tineola bisselliella*, the webbing clothes moth, is an economically important, globally distributed synanthropic pest species and member of the basal moth lineage Tineidae. *Tineola bisselliella* is facultatively keratinophagous. Therefore, their larvae can cause extensive damage, particularly to clothing, textiles, and museum specimens. Despite the economic and phylogenetic importance of *Tineola bisselliella*, there is a lack of quality genomic resources for this species, or for others within the Tineidae family. The *Tineola bisselliella* genome presented here consists of 30 pseudochromosomes (29 autosomes and 1 Z chromosome) produced using synteny alignment to a closely related species, *Tinea pellionella*. The resulting final pseudochromosome-level assembly is 243.630 Mb and has an N50 length of 8.708 Mb. The assembly is highly contiguous and has similar or improved quality compared to other available Tineidae genomes, with 93.1% of lepidopteran orthologs complete and present. Annotation of the pseudochromosome-level genome assembly with the transcriptome we produced ultimately yielded 11,267 annotated genes. Synteny alignments between the *Tineola bisselliella* genome assembly and other Tineidae genomes found evidence for numerous small rearrangements with high synteny conservation. In contrast, synteny alignments performed between *Tineola bisselliella* and the more distantly related *Bombyx mori* and *Melitea cinxia* revealed more frequent small and large rearrangements as predicted by their evolutionary divergence. The reference quality annotated genome for *Tineola bisselliella* presented here will advance our understanding of the evolution of the lepidopteran karyotype by providing a chromosome-level genome for this basal moth lineage and provide future insights into the mechanisms underlying keratin digestion in *Tineola bisselliella*.

**Significance Statement:** *Tineola bisselliella*, the webbing clothes moth, is a globally distributed synanthropic pest species that feeds on both keratin-based materials and detritus. This dietary habit can cause substantial damage to clothing, textiles, rugs, taxidermy, and museum specimens. In fact, keratinophagous organisms cause an estimated $1 billion worth of damage annually in the United States. However, the lack of a high-quality annotated reference genome for this basal moth species has thus far limited our understanding of the mechanisms underlying keratin digestion relevant to pest control efforts and has hampered efforts to investigate the broader phylogenetic relationships the Tineidae family necessary to reveal patterns of lepidopteran chromosome evolution. Here, we present the first reference quality genome for *Tineola bisselliella* and the first annotated genome for any member of the Tineidae family, providing a critical resource for the lepidopteran genomics and pest management communities.

## Introduction

Synanthropic organisms are undomesticated species that live with and exploit humans (Klegarth, 2017). The webbing clothes moth, *Tineola bisselliella*, is a synanthropic pest species with a worldwide distribution (Plarre & Krüger-Carstensen, 2011). *Tineola bisselliella* is a facultative keratinophagous species, because as larvae they can feed on both keratin and detritus (Plarre & Krüger-Carstensen, 2011; Querner, 2016). Additionally, *Tineola bisselliella* is a member of the basal lepidopteran family Tineidae, a large and diverse group of moths whose ancestral diet consisted of fungus (Davis & Gaden, 1999; Regier et al., 2015). However, many moths within Tineidae have also evolved the ability to digest keratin (Regier et al., 2015). Despite the estimated $1 billion USD worth of damage caused by keratinophagous organisms each year, research into the mechanism of keratin digestion in Tineidae moths has been greatly limited by a lack of quality annotated genomic resources (Clark et al., 2016; Metcalf & Luckmann, 1994).

Previous efforts to generate a *de novo* reference genome for *Tineola bisselliella* have resulted in fragmented contig-level genomes that fall short of the reference quality necessary to understand chromosome evolution and identify the mechanisms responsible for keratin digestion (GenBank accession no. GCA_026546545.1 and GCA_028551675.1; Clark et al., 2016). Additionally, the absence of high-quality transcriptomic data has hampered any attempts at genome annotation in the family. Therefore, additional efforts are needed to improve the existing genomic resources for this system and to advance our understanding of lepidopteran genome evolution more broadly.

A well-annotated genome is also needed to provide a foundation for investigations of the mechanisms underlying diet evolution and keratin digestion in *Tineola bisselliella*. Previous research has identified two *Bacillus* species in the gut of *T. bisselliella* larvae that express keratinases, which are hypothesized to be partially responsible for keratin digestion (Vilcinskas et al., 2020). However, the expression of serine protease or cysteine synthase genes in the moth’s genome has been proposed as an alternative mechanism for keratin digestion (Hughes & Vogler, 2006; Schwabe et al., 2021). This latter hypothesis is based on the ability of these enzymes to break the complex disulfide bonds contained in keratin (Hughes & Vogler, 2006; Schwabe et al., 2021). Unfortunately, both avenues of investigation have been limited by the lack of a quality assembled and annotated genome which has led to a lack of understanding surrounding the evolution of keratinophagy in this family.

*Tineola bisselliella* is also significant in the context of the lepidopteran phylogeny, as a member of the basal moth family Tineidae. Generating a high-quality genome for this species may shed light into lepidopteran karyotype evolution, which is of particular interest because of the large diversity in the haploid chromosome number in the group, ranging from 5 to 223 (de Vos et al., 2020). This diversity stems from the fact that Lepidoptera possess holocentric chromosomes, which have kinetochores that span the length of their chromosomes allowing chromosome fragments to be inherited (Mandrioli & Manicardi, 2020; Wright et al., 2024). These types of karyotype changes can create pre-zygotic and post-zygotic barriers to gene flow that can ultimately lead to speciation (de Vos et al., 2020). Therefore, assessment of synteny conservation between lepidopteran genomes can provide insight into karyotype changes acquired during the process of speciation.

Here, we present the first reference quality genome for *Tineola bisselliella* and the first annotated genome for any member of the Tineidae family to aid in these future research directions. Additionally, we performed preliminary synteny analyses between our pseudochromosome-level *Tineola bisselliella* genome assembly and available genomes of Tineidae family members, along with *Bombyx mori* and *Melitaea cinxia*, to investigate lepidopteran karyotype evolution.

## Results and Discussion

Whole genome sequencing of two *Tineola bisselliella* adults using PacBio HiFi technology yielded 11 Gb of HiFi data with the two libraries containing 646,360 and 355,368 reads, which were pooled to produce the final genome assembly with an estimated base coverage of 43X. Improved Phased Assembly (IPA) produced a genome assembly with 256 contigs (Figure 1A). Synteny alignment of the contig-level genome to the existing *Tinea pellionella* reference genome by Satsuma produced an assembly with a total of 89 superscaffolds, an N50 scaffold length of 5.904 Mb, and a total genome size of 243.595 Mb (Figure 1A). Satsuma then produced the final pseudochromosome assembly which contained 30 pseudochromosomes (29 autosomes and 1 Z chromosome). The N50 pseudochromosome length was 8.708 Mb and the total genome size was 243.630 Mb (Figure 1A). Additional summary statistics for the genome assembly are presented in Figure 1A.

**Figure 1.**
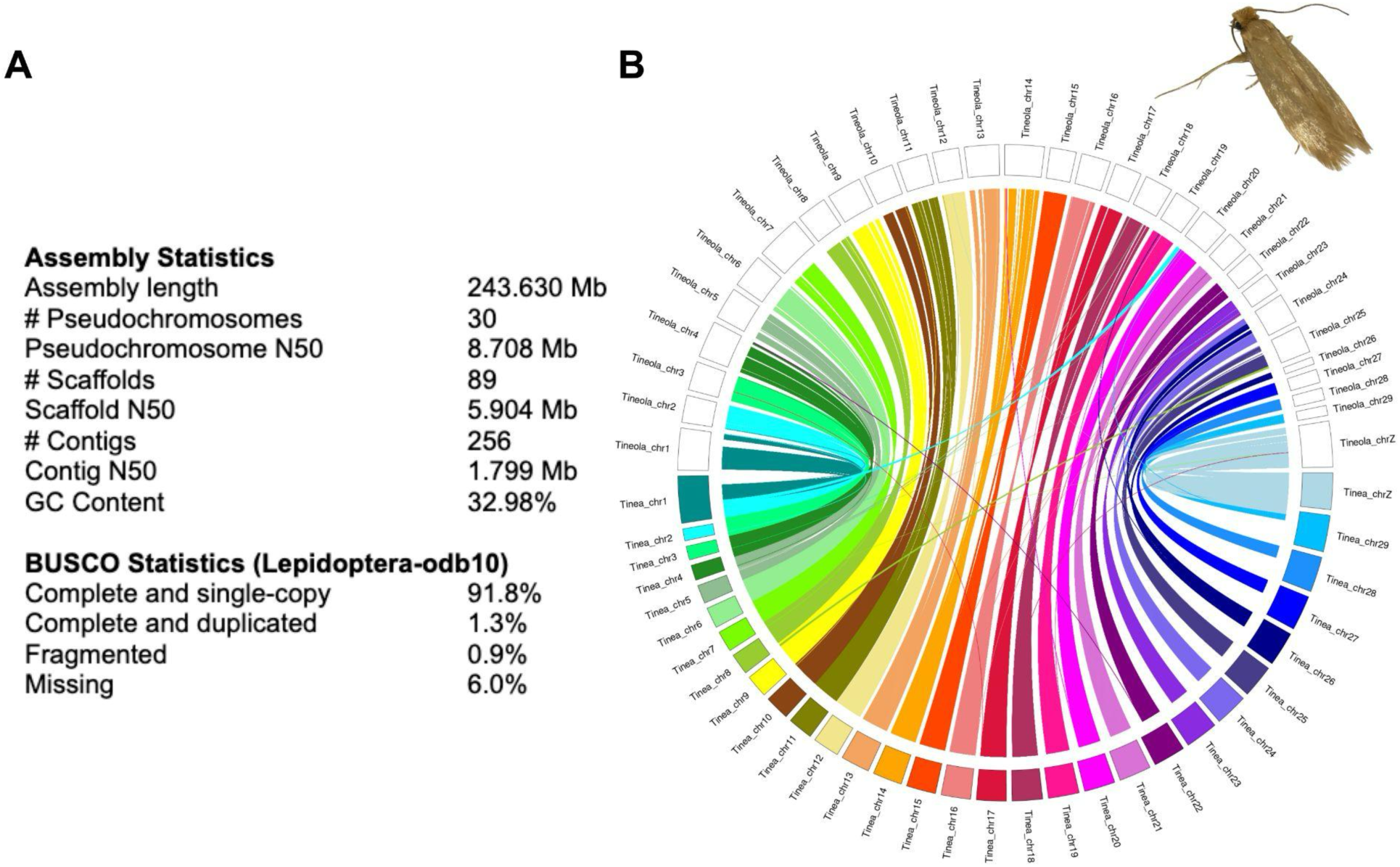
Genome assembly statistics and Circos plot of synteny alignment between *Tineola bisselliella* and *Tinea pellionella*. **A)** Assembly (top) and BUSCO (bottom) statistics of the final pseudochromosome-level *Tineola bisselliella* genome assembly. BUSCO statistics indicate percentage of orthologs complete, fragmented, or missing from the Lepidoptera-odb10 lineage dataset. **B)** Synteny alignment between the pseudochromosome-level genome assembly of *Tineola bisselliella* and the chromosome-level genome assembly of *Tinea pellionella* (GenBank accession no. GCA_948150575.1), which was used to assemble the scaffolds of the *Tineola bisselliella* genome into chromosomes by synteny-based assembly. Inset is an image of a *Tineola bisselliella* adult.

Benchmarking Universal Single-Copy Orthologs (BUSCO) analysis revealed that the quality of the *Tineola bisselliella* pseudochromosome assembly was consistent with the quality of the existing *Tinea pellionella* chromosome-level genome assembly that was produced from long read sequencing and Hi-C data (GenBank accession no. GCA_948150575.1; Figure 1A; Figure S1-S2; Table S1). Both genomes had 93% of BUSCO orthologs from the Lepidoptera-odb10 lineage dataset present as complete orthologs (Figure 1A; Figures S1-S2; Table S1). This is greater than the average quality of lepidopteran genomes available on LepBase and NCBI GenBank (81.2% BUSCO score, scaffold N50 of 1.706 Mb) reported by Ellis et al., (2021). BUSCO analysis using the Bacteria-odb9 lineage dataset identified bacterial contamination on four scaffolds of the pseudochromosome assembly, which was masked from the final assembly using BEDtools. A list of the masked regions is included in Table S2.

To aid in genome annotation, RNA sequencing data was obtained from *Tineola bisselliella* adults (328,802,057 paired-end reads), late-stage larvae (335,986,188 paired-end reads), and early-stage larvae (302,067,521 paired-end reads). After quality filtering, adapter trimming, polyG tail trimming, and assembly of contigs using Trinity, the transcriptome assembly contained 231,316 contigs. After discarding contigs that were shorter than 500 bp, 99,930 contigs remained. The final transcriptome assembly contained 58,509 transcripts, which consisted of 39,404 genes. EggNOG-mapper v2 functionally annotated 13,615 genes in the transcriptome. Genome annotation using the MAKER pipeline identified 11,267 protein coding genes corresponding to 13,114 mRNA transcripts, and 94% of these annotations had an annotation edit distance (AED) score of 0.5 or less after the second round of *ab initio* gene prediction (Table 1; Figure S4). 10,769 of the identified protein coding genes were functionally annotated by blastp (Table 1).

**Table 1.**
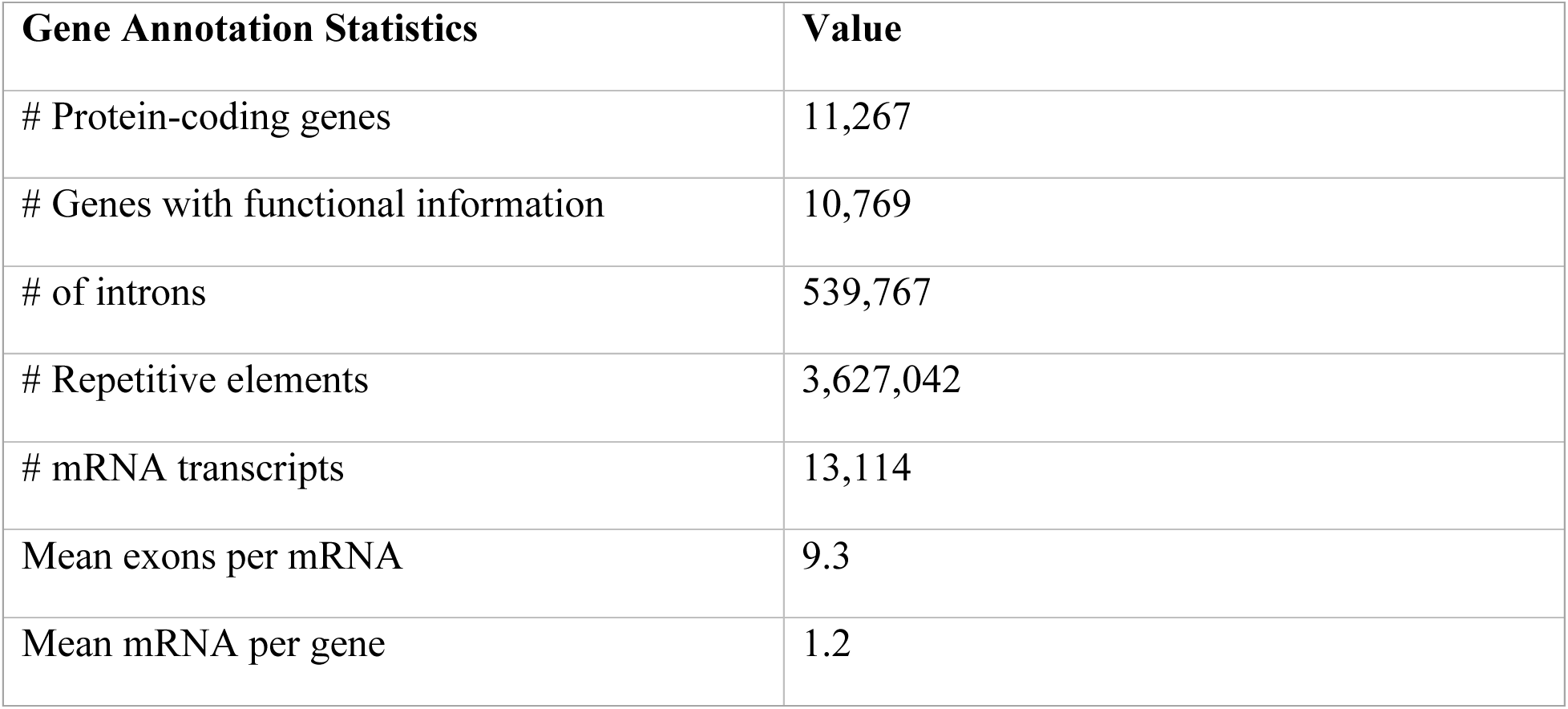
Summary statistics for the final *Tineola bisselliella* pseudochromosome genome annotation.

Interestingly, our *Tineola bisselliella* genome includes annotations with transcriptomic evidence for both serine proteases and cysteine synthases. This is consistent with a previously published analysis of the *Tineola bisselliella* larval gut, which found evidence for the expression of both serine proteases or cysteine synthases in the larval gut of *Tineola bisselliella* without a bacterial source (Hughes & Vogler, 2006; Schwabe et al., 2021). Taken together, these results suggest that the ability to digest keratin is encoded within the *Tineola bisselliella* genome. However, serine proteases are found in the gut of all Lepidoptera and function as a primary step in plant protein digestion (Srinivasan et al., 2006). This suggests that the expression of serine proteases alone may not be sufficient to have facilitated keratin digestion, because of the close relatedness of serine proteases in the *Tineola bisselliella* gut to the serine proteases in other Lepidoptera (Hughes & Vogler, 2006). Nevertheless, an alternative hypothesis is that the digestion of keratin is facilitated by a combination of symbionts in the gut microbiome, enzymes encoded in the genome, and the low oxidation-reduction potential of the larval gut first observed by Waterhouse (1952). Future work investigating this possibility is needed, and the annotation resource presented here will be essential to understanding the role of these candidate enzymes in keratin digestion.

Our assembly and annotation also provide an important resource for investigating the history of lepidopteran chromosome evolution. Chromosomal rearrangements are expected in lepidopteran genomes because their chromosomes are holocentric and therefore able to tolerate inheritance of chromosome fission and fusion events (Mandrioli & Manicardi, 2020; Wright et al., 2024). Synteny alignment of the *Tineola bisselliella* and *Tinea pellionella* genomes demonstrated evidence for high synteny conservation between the two species (GenBank accession no. GCA_948150575.1; Figure 1B). However, the alignment also illustrated that small chromosomal rearrangements between the two species have occurred and, therefore, we retained these structural differences during the synteny-based assembly of the *Tineola bisselliella* genome into pseudochromosomes (Figure 1B). Additional synteny alignments of the *Tineola bisselliella* pseudochromosome genome assembly with two other Tineidae family genomes, *Monopis laevigella* and *Tinea trinotella*, displayed similar patterns of high synteny conservation with small rearrangements (GenBank accession no. GCA_947458855.1 and GCA_905220615.1; Figures S5-S6).

In contrast to these patterns of high synteny conservation among Tineids, synteny alignments of the *Tineola bisselliella* pseudochromosome genome assembly to more distantly-related species, *Bombyx mori* and *Melitaea cinxia*, displayed evidence for frequent chromosomal rearrangements of both large and small genome regions (GenBank accession no. GCA_014905235.2 and GCA_905220565.1; Figures S5-S6). This is not unexpected, because the Tineidae family is a basal clade in relation to Bombycoidea and Papilionoidea (Kawahara et al., 2019). Importantly, the synteny alignment of *Tineola bisselliella* to *Melitaea cinxia* provided unique insights into the evolution of the lepidopteran ancestral karyotype, as *Melitaea cinxia* is believed to be the only butterfly species that has retained the ancestral karyotype of 31 (30 autosomes) (Ahola et al., 2014; de Vos et al., 2020). Reduction in chromosome number from the ancestral karyotype is hypothesized to be the result of whole chromosomal fusions, where two independent chromosomes fuse to make one super chromosome (Ahola et al., 2014). However, when we performed a synteny alignment of the *Tineola bisselliella* pseudochromosome genome assembly to *Melitaea cinxia,* our assembly contained no whole chromosomal fusions of *Melitaea cinxia* chromosomes. Instead, there were many small fragmentations that fused with existing ancestral chromosomes. Therefore, our findings are consistent with a scenario in which the reduction in karyotype size in *Tineola bisselliella* is due to many small chromosomal fragmentation and fusion events, and not whole ancestral chromosome fusion events. However, future Hi-C confirmation of the *Tineola bisselliella* pseudochromosome genome will be needed to confirm this hypothesis.

In summary, the pseudochromosome-level assembly generated here is the first reference quality genome for *Tineola bisselliella*. It also represents the first annotated genome in the Tineidae family, and will serve as an important genomic resource for the scientific community. Finally, we believe that it will help improve our knowledge of basal moth taxonomy, contribute to our understanding of the evolutionary mechanisms driving patterns of speciation within the Tineidae family, and serve as an important resource for future investigation of the evolution of keratin digestion.

## Materials and Methods

### Sample Acquisition, DNA and RNA Extraction, and Sequencing

Two *Tineola bisselliella* larvae were obtained from an infested fur acquired from a museum and were sent to the University of Delaware DNA Sequencing and Genotyping Center for high molecular weight DNA extraction and long-read whole genome sequencing in March 2021. The two larvae were individually barcoded and sequenced in a PacBio SMRT cell at the University of Delaware producing 11 Gb of HiFi long-reads.

Ten live *T. bisselliella* individuals (five early-stage larvae, three late-stage larvae, and two adults) were sampled from a lab colony established in June 2022. RNA was extracted using a Qiagen RNeasy Micro Kit following manufacturer’s instructions, except samples were homogenized with lysis buffer and glass beads for 1 minute prior to extraction. RNA quality was assessed visually by performing agarose gel electrophoresis and quantified using NanoDrop. The highest quality extractions, specifically two early-stage larvae, three late-stage larvae, and two adults, were sent to the Tufts University Genomics Core Facility for library preparation and sequencing in August 2023. The samples were separated into three barcoded pools based on developmental stage (*i.e.,* early-stage larvae, late-stage larvae, or adult) and were then prepped using the Illumina Stranded mRNA Prep method. Paired-end sequencing of all three libraries was performed on an Illumina NovaSeq Flowcell.

### Genome Assembly

The University of Delaware performed demultiplexing, quality filtering, and Improved Phased Assembly (IPA) of the reads (GitHub: https://github.com/PacificBiosciences/pbipa), yielding a contig-level assembly. This genome assembly was then mapped by synteny to a chromosome-level assembly of *Tinea pellionella* (GenBank accession no. GCA_948150575.1), using Satsuma v.3.0 Chromosembler (Grabherr et al., 2010). Satsuma first produced an intermediate scaffold-level assembly, which was mapped again to the *Tinea pellionella* genome assembly to produce the final pseudochromosome-level assembly. Genome assembly statistics for all assemblies were computed using BBMap’s stats.sh script (Bushnell, 2014). To assess the completeness of the final pseudochromosome-level assembled genome, Benchmarking Universal Single-Copy Orthologs (BUSCO) v3.0.2 was performed using the BUSCO v4 Lepidoptera-odb10 (Simao et al., 2015; Manni et al., 2021). Reference quality genomes for *Bombyx mori* (GenBank accession no. GCA_014905235.2), *Tinea pellionella* (GenBank accession no. GCA_948150575.1), and *Monopis laevigella* (GenBank accession no. GCA_947458855.1) were obtained from NCBI GenBank, and each of these genomes were also assessed using BUSCO for genome quality comparison. Bacterial contamination was assessed using BUSCO (v3.0.2) with the Bacteria-odb9 lineage set (Simao et al., 2015). Bacterial contamination was hard-masked using BEDtools (Quinlan and Hall, 2010).

### Transcriptome Assembly and Annotation

The paired-end RNAseq data were assembled and annotated following methods presented in Rivera and Davies (2021). The assembled transcriptome was submitted to the eggNOG-mapper v2 online platform and annotated using the eggNOG 5.0 dataset (Cantalapiedra et al., 2021; Huerta-Cepas et al., 2019). Gene Ontology (GO) annotations along with gene names were extracted from the eggNOG-mapper v2 output and used to produce an annotated FASTA file.

### Genome Annotation

The pseudochromosome genome assembly was annotated following the MAKER pipeline detailed in Cantarel et al. (2008) using MAKER v3.01.04, with the modifications detailed in Figure S3. All available EST evidence for the Tineidae family was obtained from NCBI GenBank in November 2023 (Clark et al., 2016) to incorporate as evidence in addition to the generated transcriptome. Protein evidence was gathered from the November 2022 UniProt Lepidoptera release and eggNOG-mapper v2 annotation of the genome assembly (The UniProt Consortium, 2023; Cantalapiedra et al., 2021; Huerta-Cepas et al., 2019). Repeat evidence was obtained from GIRI RepBase’s invertebrate repeat November 2022 release (Bao et al., 2015). All evidence was aligned to the genome assembly using RepeatMasker, Exonerate, and BLAST+ (Bao et al., 2015; Camacho et al., 2009; Slater & Birney, 2005).

The *ab initio* gene prediction software SNAP and Augustus (Korf, 2004; Stanke et al., 2006) were trained using hidden Markov models (HMM) generated by BEDtools and BUSCO (Quinlan & Hall, 2010; Manni et al., 2021). After each round of *ab initio* gene prediction, the number of gene models and the corresponding AED scores were recorded and plotted to evaluate the effectiveness of the prediction software (Figure S4). Training and running of *ab initio* gene prediction was repeated for a total of two rounds. blastp was used to functionally annotate protein-coding genes (Camacho et al., 2009). Summary statistics of the genome annotation were then compiled using the agat_sp_statistics.pl script included in AGAT (GitHub: https://github.com/NBISweden/AGAT).

### Synteny Alignments and Visualization

The *Tineola bisselliella* pseudochromosome-level assembly was aligned to all three available Tineidae chromosome-level genome assemblies: *Tinea pellionella* (GenBank accession no. GCA_948150575.1), *Tinea trinotella* (GenBank accession no. GCA_905220615.1), and *Monopis laevigella* (GenBank accession no. GCA_947458855.1), in addition to a *Bombyx mori* genome assembly (GenBank accession no. GCA_014905235.2) and a *Melitaea cinxia* genome assembly (GenBank accession no. GCA_905220565.1). Satsuma was used to perform the synteny analysis (Grabherr et al., 2010), and the R packages RIdeogram and Circlize were used to visualize the outputs of Satsuma (Hao et al., 2020; Gu et al., 2014).

## Data Availability

The raw reads from PacBio HiFi genome sequencing and Illumina raw transcriptome short reads are deposited under BioProject PRJNA1102366. The pseudochromosome genome assembly has been submitted to NCBI (JBFQDO000000000). Upon acceptance of this manuscript the genome assembly and annotation will be available on Dryad. All code associated with this project can be found in the following GitHub Repository: https://github.com/jasalq/Tineola_bisselliella_genome.

## Supplementary Materials

**Table S1.**
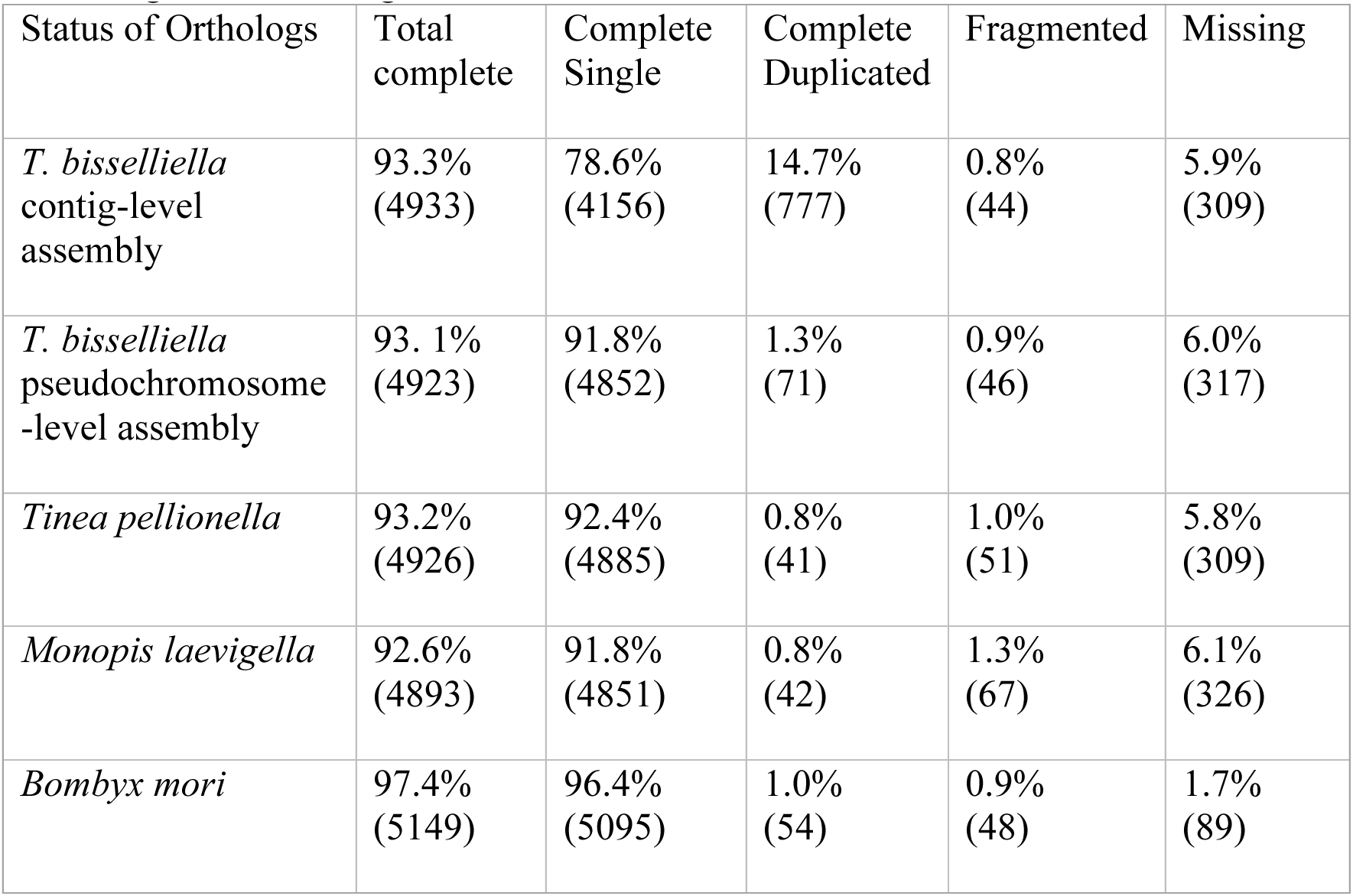
Results of BUSCO Analysis. Percent and number (in parentheses) of orthologs found in each assembly determined from BUSCO analysis of the genome assemblies using the Lepidoptera-odb10 lineage dataset containing 5,286 orthologs.

**Table S2.**
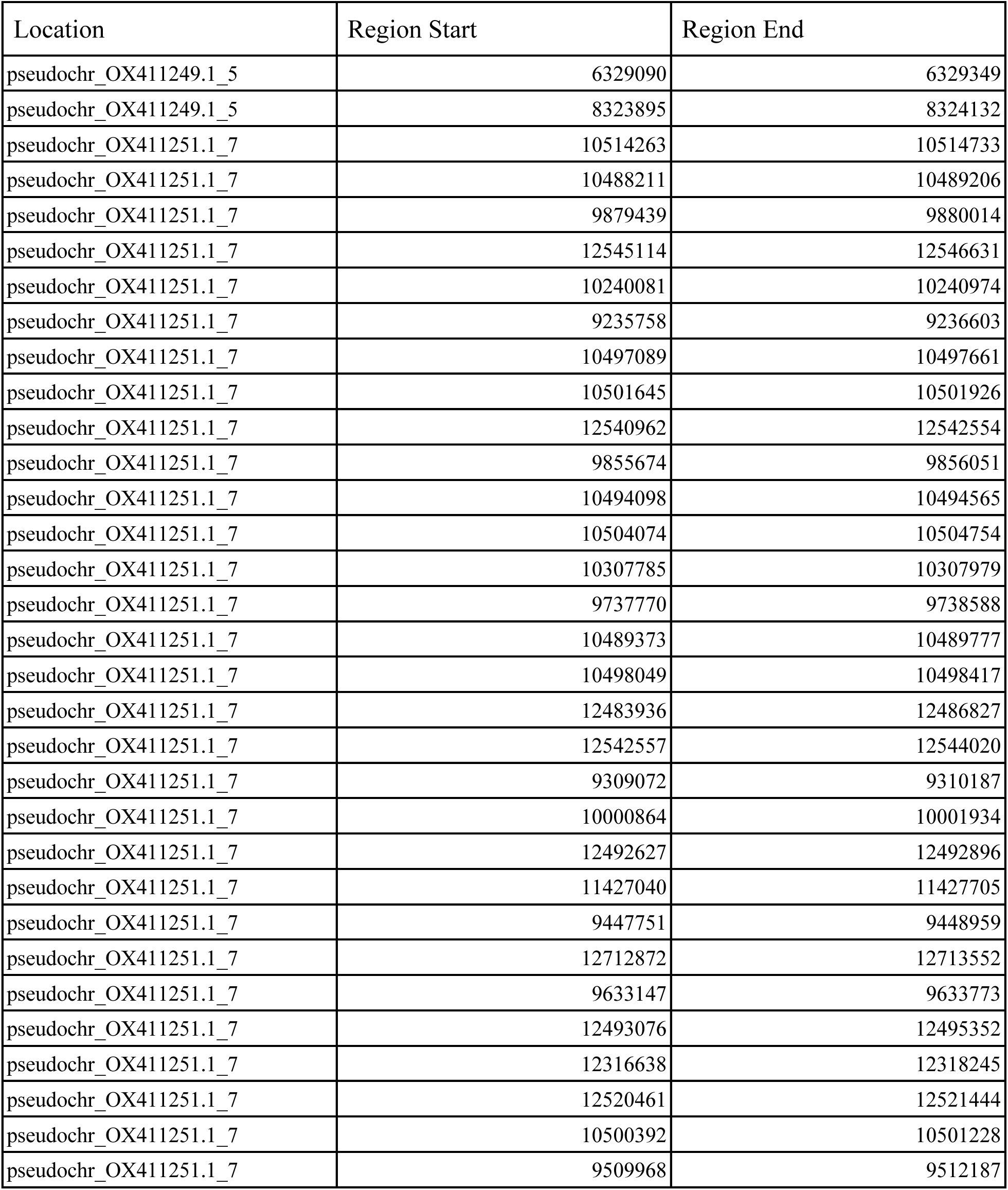
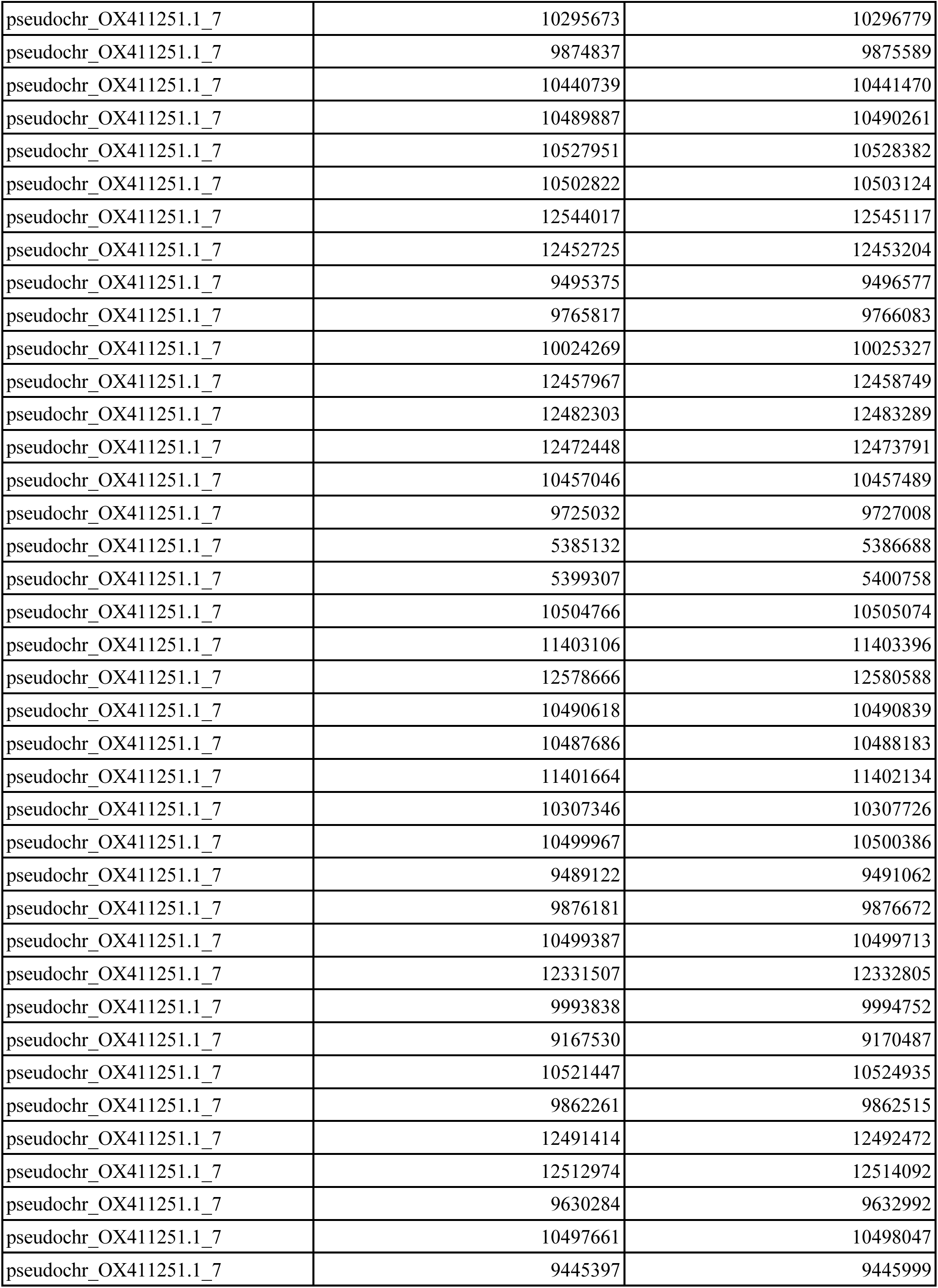
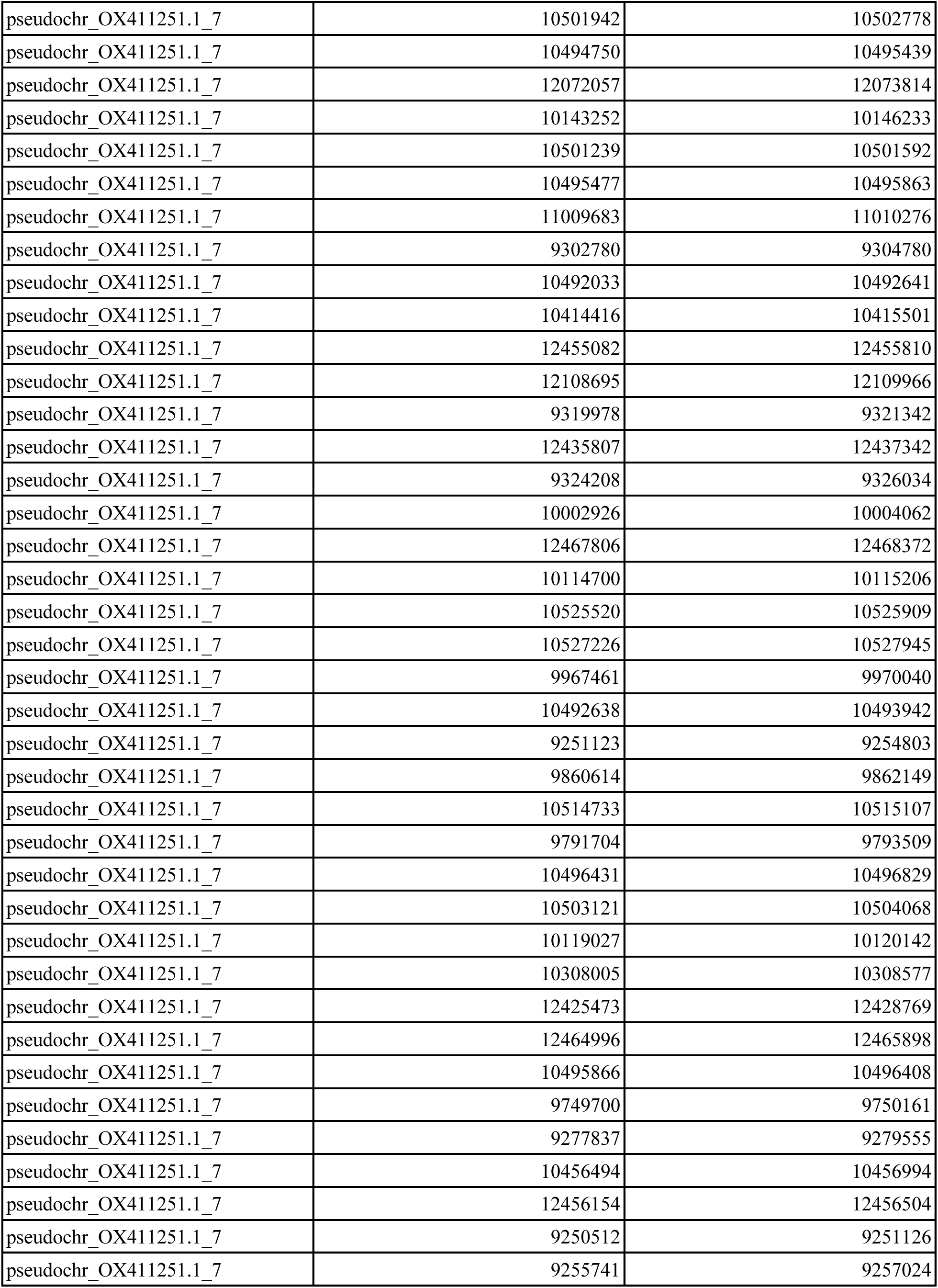
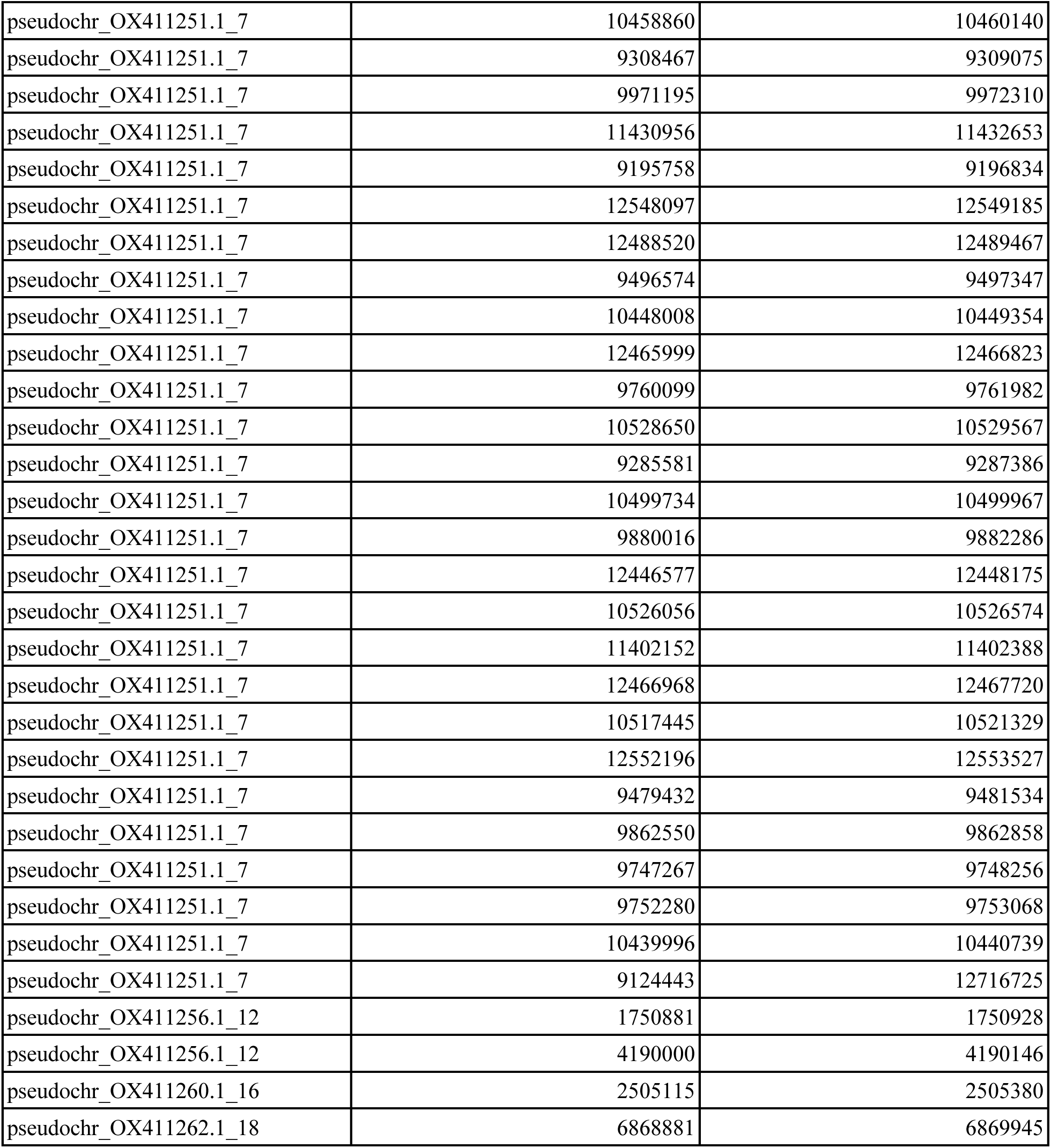
Regions of Genome Masked Due to Bacterial Contamination. Bacterial contamination was found using BUSCO analysis using the Bacteria-odb9 lineage. Contamination was masked from the final assembly using BEDtools. This table presents the coordinates at the beginning (‘Region Start’) and end (‘Region End’) of the bacterial contamination regions masked in the final assembly.

**Figure S1.**
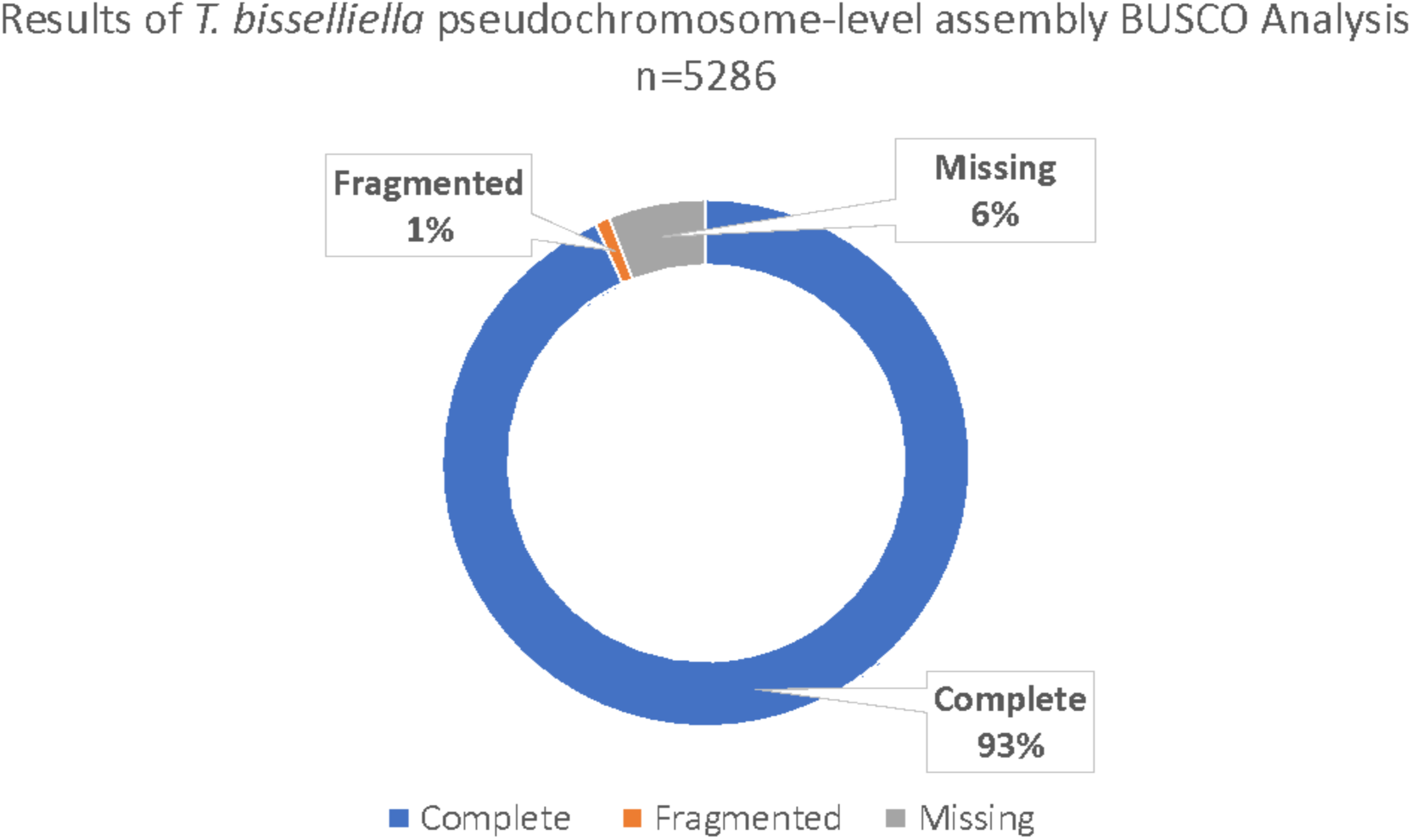
BUSCO results of *Tineola bisselliella* pseudochromosome-level assembly. Results from the BUSCO analysis of the pseudochromosome-level assembly including the combined percentage of complete orthologs (single copy and duplicated), as well as the percent of fragmented and missing orthologs. The exact number of orthologs in each category can be seen in Table S1.

**Figure S2.**
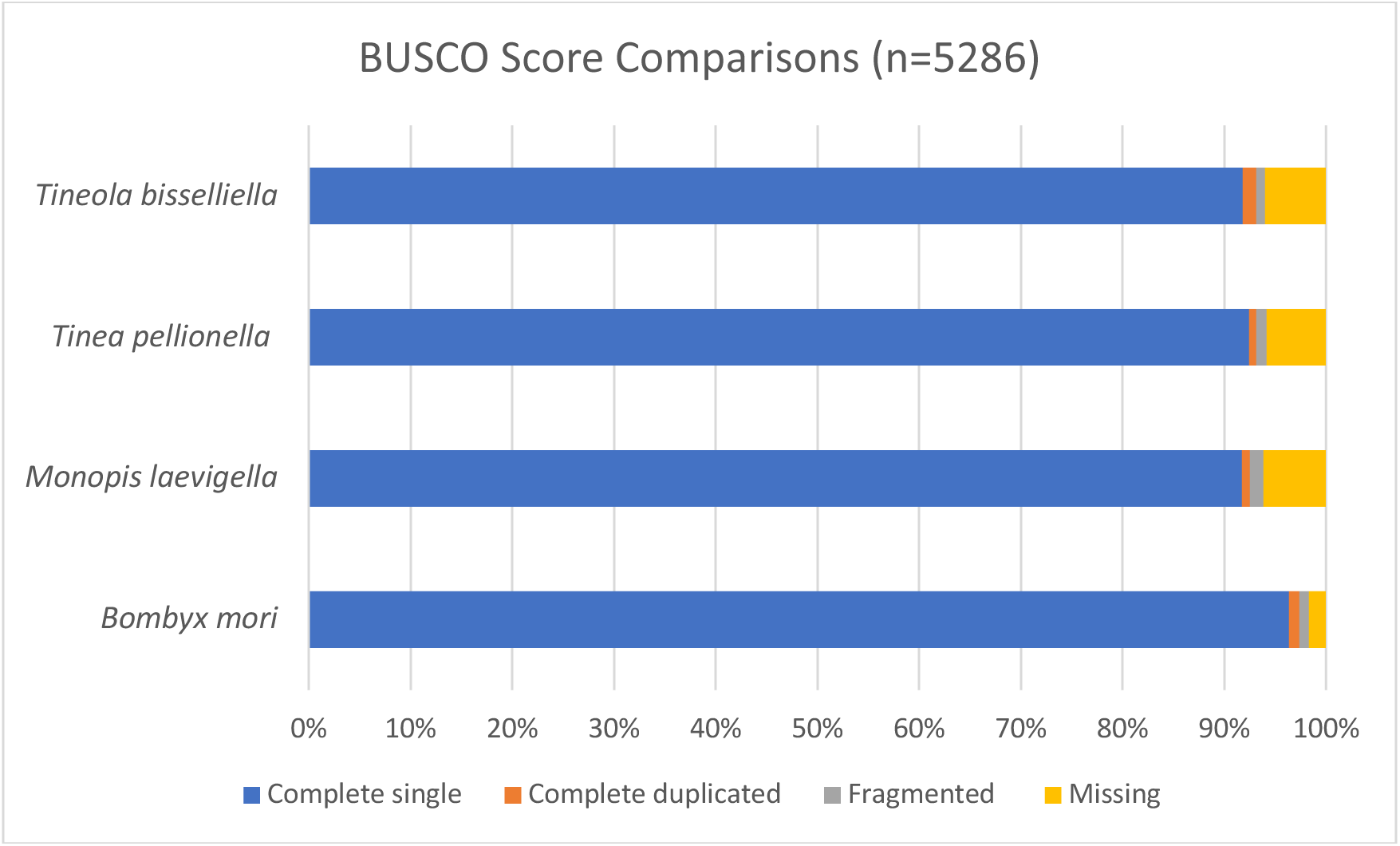
Comparison of BUSCO results of different genome assemblies with *Tineola bisselliella* pseudochromosome-level assembly. Percent of complete single copy, complete duplicated, fragmented, and missing orthologs from Lepidoptera_odb10 lineage dataset found in our *Tineola bisselliella* pseudochromosome-level genome assembly, *Tinea pellionella* assembly (GenBank accession no. GCA_948150575.1), *Monopis laevigella* assembly (GenBank accession no. GCA_947458855.1), and *Bombyx mori* assembly GenBank accession no. GCA_014905235.2). The exact number of orthologs in each category can be found in Table S1.

**Figure S3.**
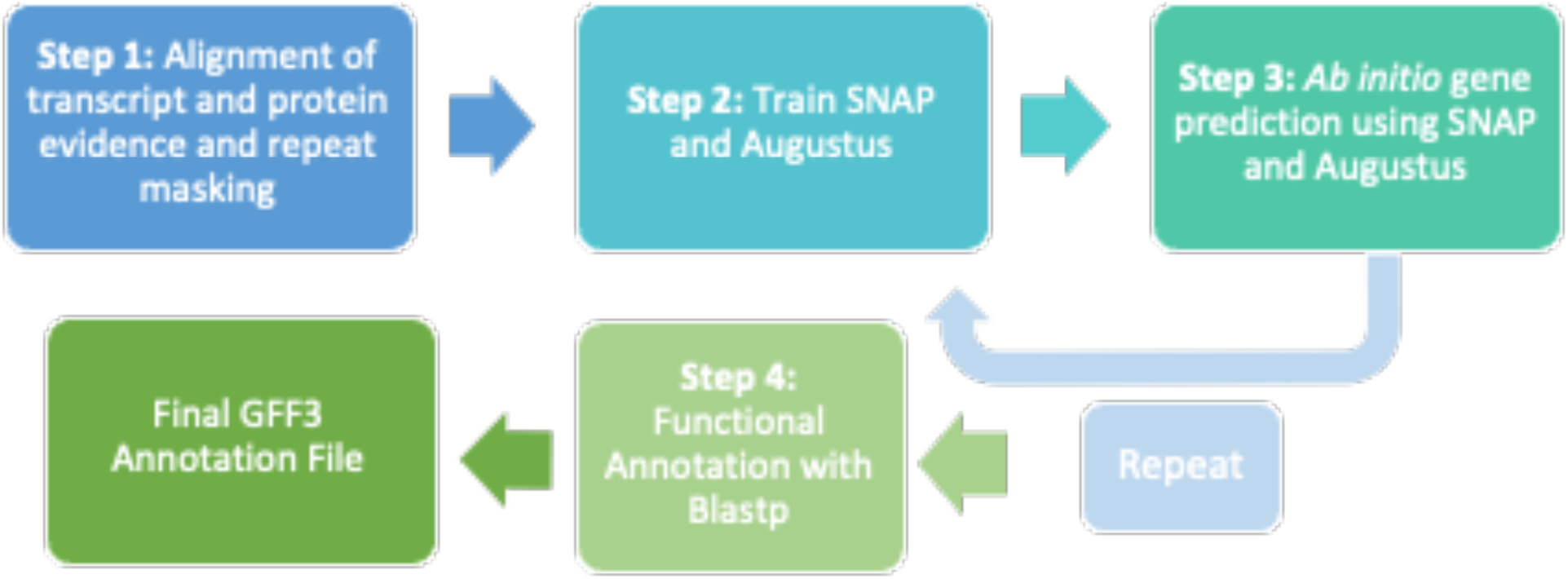
Modified MAKER genome annotation pipeline. The modified workflow for MAKER v3.01.04 genome annotation (Cantarel et al., 2008) used here. Step 1, BLAST+ was used to align transcript evidence, Exonerate was used to align protein evidence, and RepeatMasker was used to mask repetitive elements. Step 2, the *ab initio* gene prediction programs SNAP and Augustus were trained prior to each gene prediction round (step 3). The process of training and gene prediction (steps 2 and 3) was repeated until quality annotations were obtained, which took four rounds of *ab initio* gene training and prediction for the contig-level assembly and two rounds for the pseudochromosome-level assembly. During the last step (step 4), proteins were functionally annotated using blastp. This produced a final GFF3 file where all annotations were compiled.

**Figure S4.**
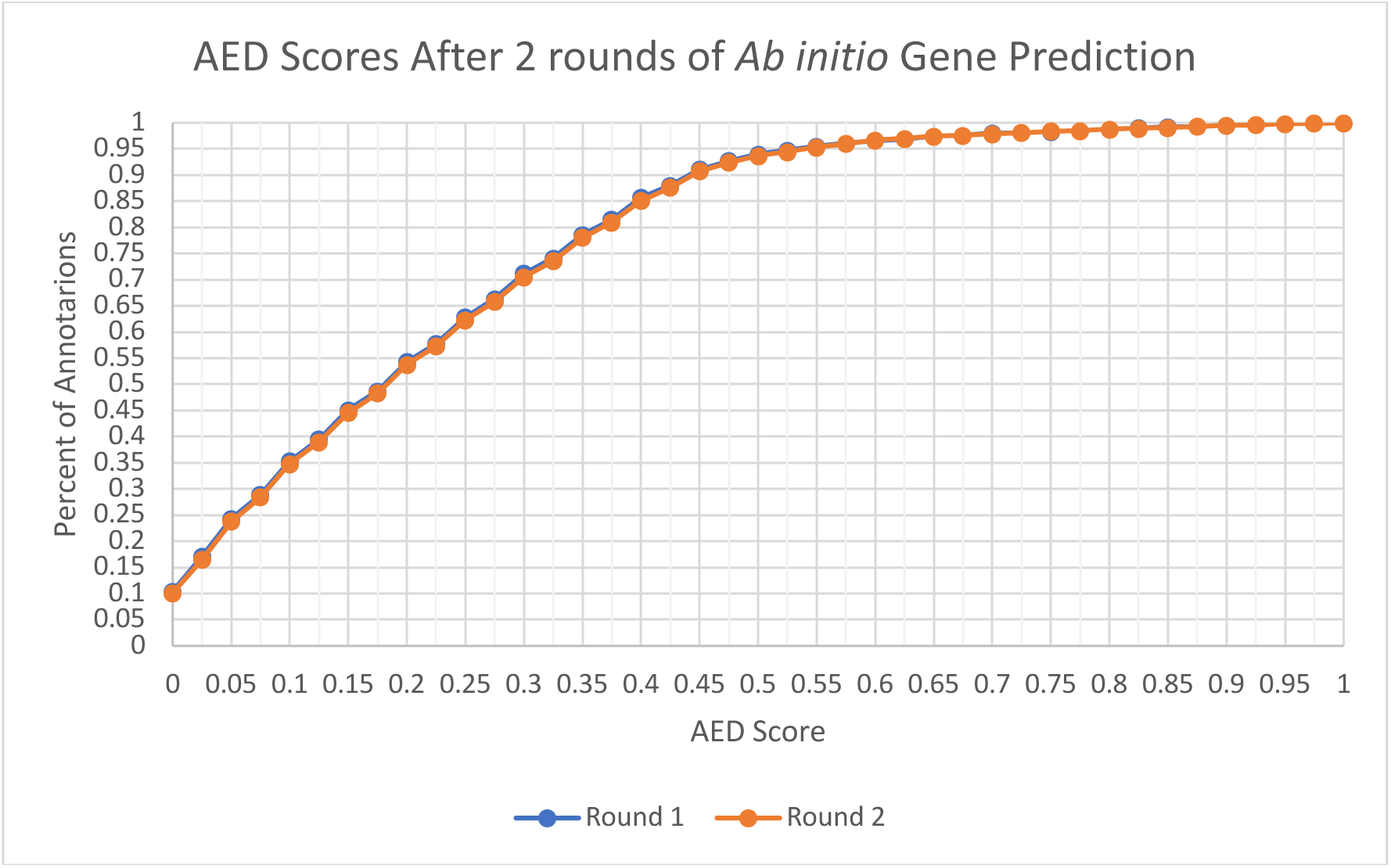
Pseudochromosome-level genome assembly Annotation Edit Distance (AED) scores after two rounds of *ab initio* gene prediction. Percent of annotations at each AED score after each round of *ab initio* gene prediction by SNAP and Augustus of the pseudochromosome-level genome assembly. AED scores have a range of 0 to 1. An AED score of 0 means the annotation is completely supported by the available evidence and an AED score of 1 means that there is no evidence supporting the annotation. The graph shows little to no difference in the percent of annotations at each AED score between the first and second round of *ab initio* gene prediction.

**Figure S5.**
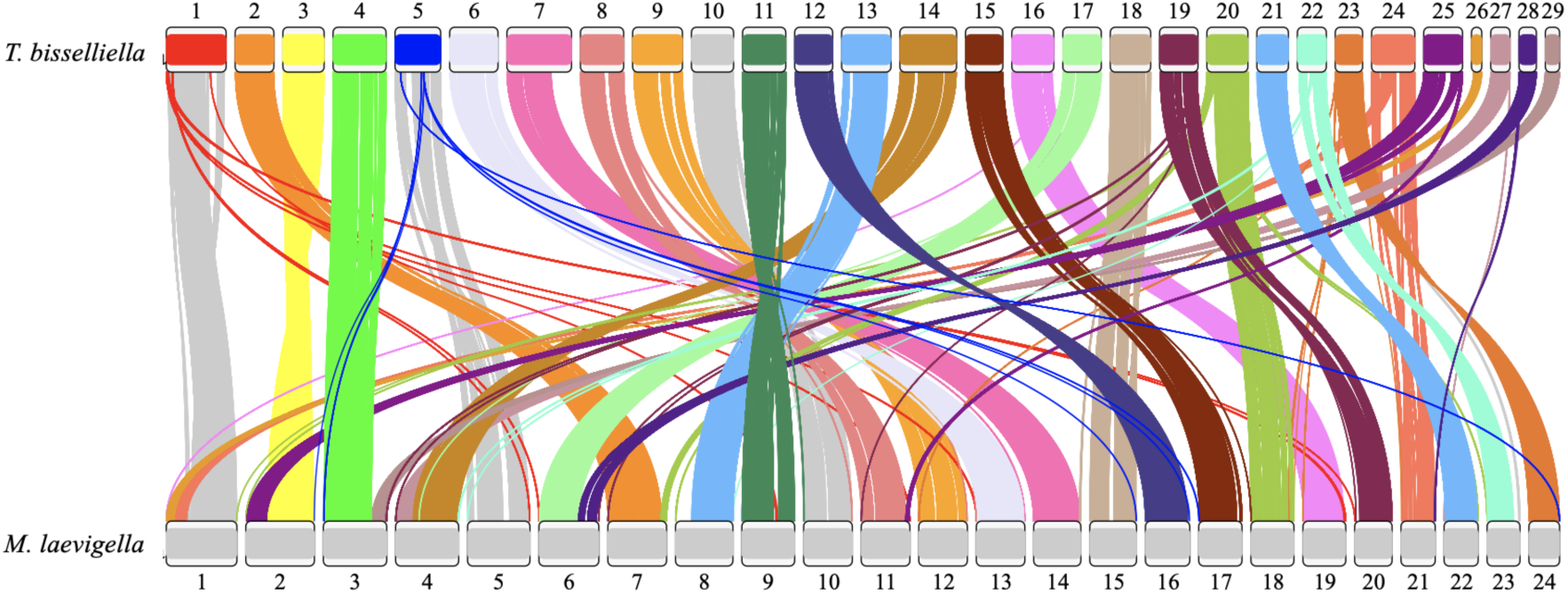
Synteny alignment of Tineola bisselliella and Monopis laevigella. Synteny alignment of *Tineola bisselliella* pseudochromosome-level genome assembly and *Monopis laevigella* genome assembly (GenBank accession no. GCA_947458855.1) produced by Satsuma. Conserved synteny is illustrated by grey lines. Chromosomal rearrangements are colored corresponding to their position in the *Tineola bisselliella* pseudochromosome genome assembly. Sex chromosomes are not included.

**Figure S6.**
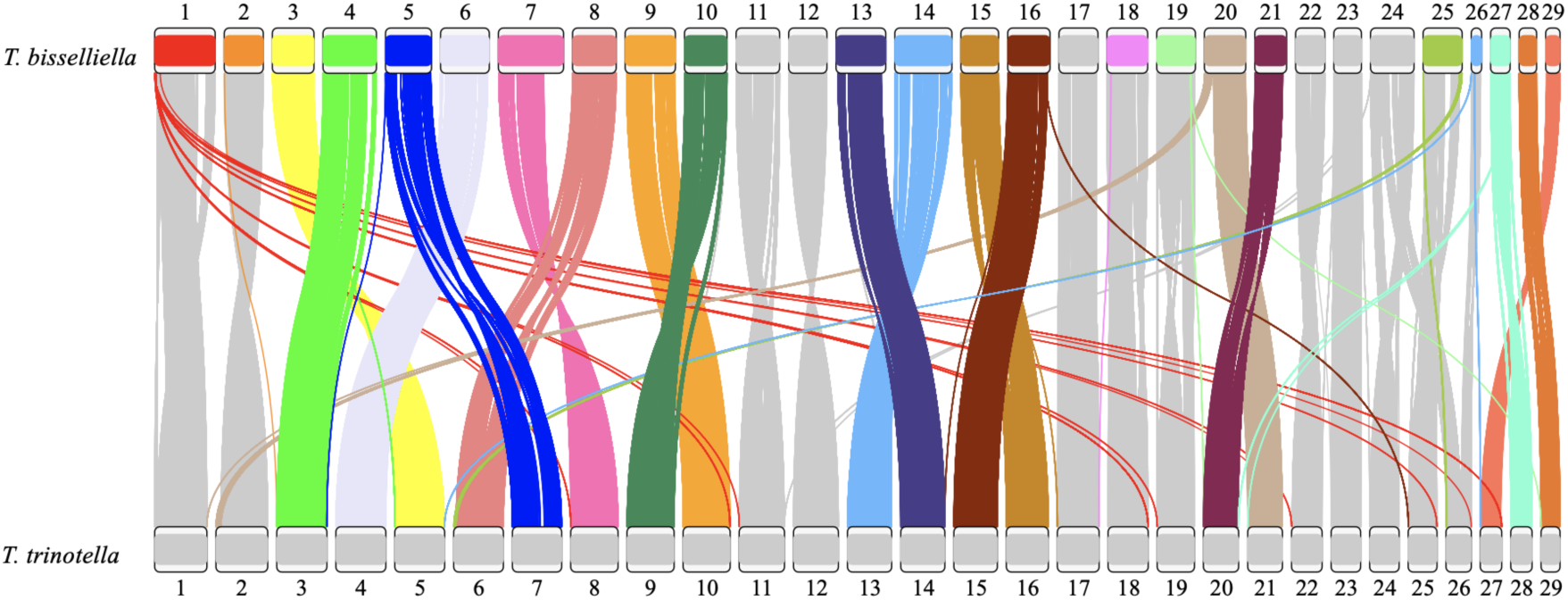
Synteny alignment of Tineola bisselliella and Tinea trinotella. Synteny alignment of *Tineola bisselliella* pseudochromosome-level genome assembly and *Tinea trinotella* genome assembly (GenBank accession no. GCA_905220615.1) produced by Satsuma. Conserved synteny is illustrated by grey lines. Chromosomal rearrangements are colored corresponding to their position in the *Tineola bisselliella* pseudochromosome genome assembly. Sex chromosomes are not included.

**Figure S7.**
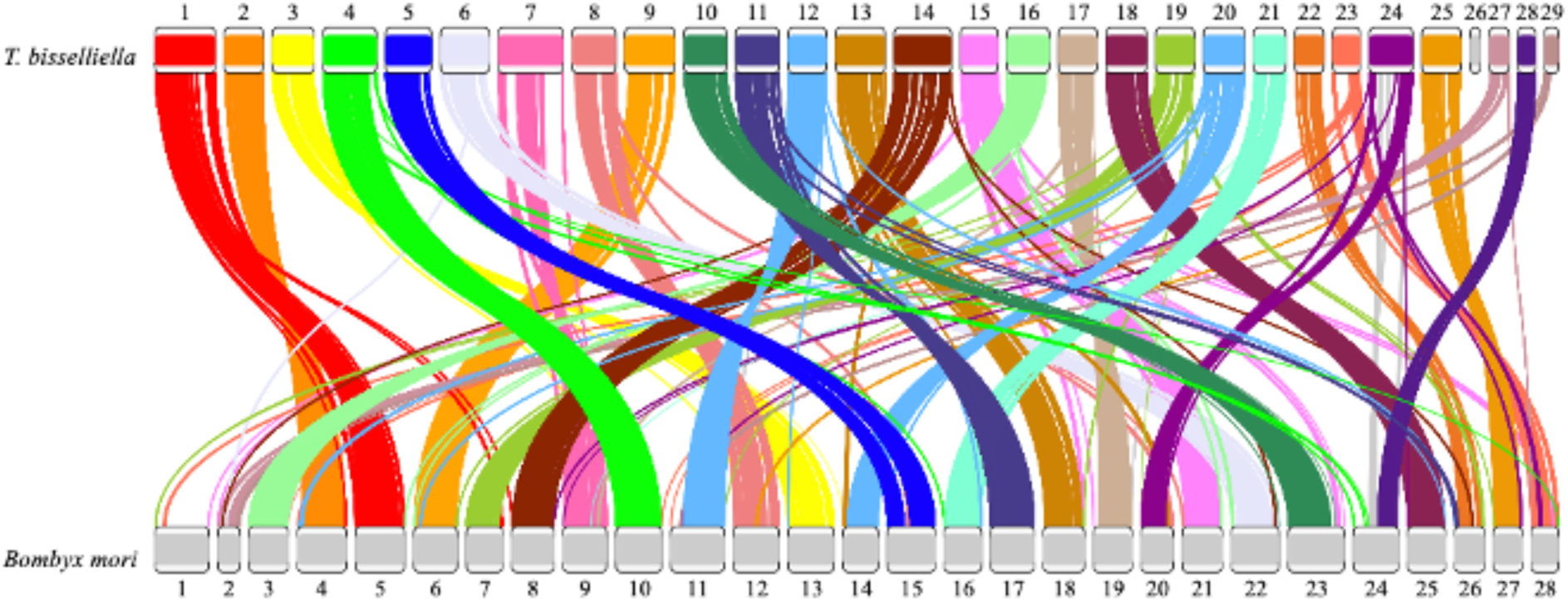
Synteny alignment of *Tineola bisselliella* and *Bombyx mori*. Synteny alignment of *Tineola bisselliella* pseudochromosome-level genome assembly and *Bombyx mori* genome assembly (GenBank accession no. GCA_014905235.2) produced by Satsuma. Conserved synteny is illustrated by grey lines. Chromosomal rearrangements are colored corresponding to their position in the *Tineola bisselliella* pseudochromosome genome assembly. Sex chromosomes are not included.

**Figure S8.**
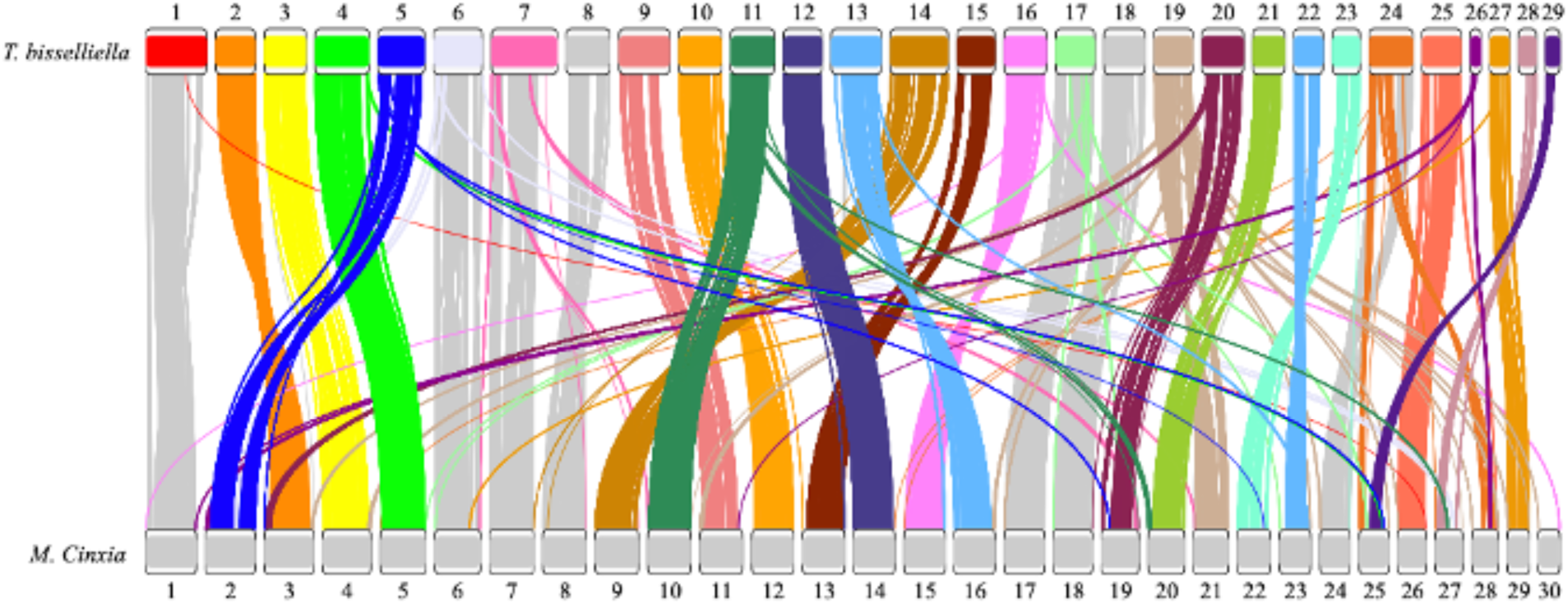
Synteny alignment of *T. bisselliella* and *Melitaea cinxia*. Synteny alignment of *Tineola bisselliella* pseudochromosome-level genome assembly and *Melitaea cinxia* genome assembly (GenBank accession no. GCA_905220565.1) produced by Satsuma. Conserved synteny is illustrated by grey lines. Chromosomal rearrangements are colored corresponding to their position in the *Tineola bisselliella* pseudochromosome genome assembly. Sex chromosomes are not included

