## Supplementary material for "De Novo Genome Assembly and Annotation for the Webbing Clothes Moth (*Tineola bisselliella*): A Globally Distributed, Economically Important Pest": Table S1

| Status of Orthologs | Total complete | Complete Single | Complete Duplicated | Fragmented | Missing |
| --- | --- | --- | --- | --- | --- |
| *T. bisselliella* contig-level assembly | 93.3%  (4933) | 78.6%  (4156) | 14.7%  (777) | 0.8%  (44) | 5.9%  (309) |
| *T. bisselliella* pseudochromosome-level assembly | 93. 1%  (4923) | 91.8%  (4852) | 1.3%  (71) | 0.9%  (46) | 6.0%  (317) |
| *Tinea pellionella* | 93.2%  (4926) | 92.4%  (4885) | 0.8%  (41) | 1.0%  (51) | 5.8%  (309) |
| *Monopis laevigella* | 92.6%  (4893) | 91.8%  (4851) | 0.8%  (42) | 1.3%  (67) | 6.1%  (326) |
| *Bombyx mori* | 97.4%  (5149) | 96.4%  (5095) | 1.0%  (54) | 0.9%  (48) | 1.7%  (89) |
