## Supplementary material for "De Novo Genome Assembly and Annotation for the Webbing Clothes Moth (*Tineola bisselliella*): A Globally Distributed, Economically Important Pest": Table S2

| Location | Region Start | Region End |
| --- | --- | --- |
| pseudochr_OX411249.1_5 | 6329090 | 6329349 |
| pseudochr_OX411249.1_5 | 8323895 | 8324132 |
| pseudochr_OX411251.1_7 | 10514263 | 10514733 |
| pseudochr_OX411251.1_7 | 10488211 | 10489206 |
| pseudochr_OX411251.1_7 | 9879439 | 9880014 |
| pseudochr_OX411251.1_7 | 12545114 | 12546631 |
| pseudochr_OX411251.1_7 | 10240081 | 10240974 |
| pseudochr_OX411251.1_7 | 9235758 | 9236603 |
| pseudochr_OX411251.1_7 | 10497089 | 10497661 |
| pseudochr_OX411251.1_7 | 10501645 | 10501926 |
| pseudochr_OX411251.1_7 | 12540962 | 12542554 |
| pseudochr_OX411251.1_7 | 9855674 | 9856051 |
| pseudochr_OX411251.1_7 | 10494098 | 10494565 |
| pseudochr_OX411251.1_7 | 10504074 | 10504754 |
| pseudochr_OX411251.1_7 | 10307785 | 10307979 |
| pseudochr_OX411251.1_7 | 9737770 | 9738588 |
| pseudochr_OX411251.1_7 | 10489373 | 10489777 |
| pseudochr_OX411251.1_7 | 10498049 | 10498417 |
| pseudochr_OX411251.1_7 | 12483936 | 12486827 |
| pseudochr_OX411251.1_7 | 12542557 | 12544020 |
| pseudochr_OX411251.1_7 | 9309072 | 9310187 |
| pseudochr_OX411251.1_7 | 10000864 | 10001934 |
| pseudochr_OX411251.1_7 | 12492627 | 12492896 |
| pseudochr_OX411251.1_7 | 11427040 | 11427705 |
| pseudochr_OX411251.1_7 | 9447751 | 9448959 |
| pseudochr_OX411251.1_7 | 12712872 | 12713552 |
| pseudochr_OX411251.1_7 | 9633147 | 9633773 |
| pseudochr_OX411251.1_7 | 12493076 | 12495352 |
| pseudochr_OX411251.1_7 | 12316638 | 12318245 |
| pseudochr_OX411251.1_7 | 12520461 | 12521444 |
| pseudochr_OX411251.1_7 | 10500392 | 10501228 |
| pseudochr_OX411251.1_7 | 9509968 | 9512187 |
| pseudochr_OX411251.1_7 | 10295673 | 10296779 |
| pseudochr_OX411251.1_7 | 9874837 | 9875589 |
| pseudochr_OX411251.1_7 | 10440739 | 10441470 |
| pseudochr_OX411251.1_7 | 10489887 | 10490261 |
| pseudochr_OX411251.1_7 | 10527951 | 10528382 |
| pseudochr_OX411251.1_7 | 10502822 | 10503124 |
| pseudochr_OX411251.1_7 | 12544017 | 12545117 |
| pseudochr_OX411251.1_7 | 12452725 | 12453204 |
| pseudochr_OX411251.1_7 | 9495375 | 9496577 |
| pseudochr_OX411251.1_7 | 9765817 | 9766083 |
| pseudochr_OX411251.1_7 | 10024269 | 10025327 |
| pseudochr_OX411251.1_7 | 12457967 | 12458749 |
| pseudochr_OX411251.1_7 | 12482303 | 12483289 |
| pseudochr_OX411251.1_7 | 12472448 | 12473791 |
| pseudochr_OX411251.1_7 | 10457046 | 10457489 |
| pseudochr_OX411251.1_7 | 9725032 | 9727008 |
| pseudochr_OX411251.1_7 | 5385132 | 5386688 |
| pseudochr_OX411251.1_7 | 5399307 | 5400758 |
| pseudochr_OX411251.1_7 | 10504766 | 10505074 |
| pseudochr_OX411251.1_7 | 11403106 | 11403396 |
| pseudochr_OX411251.1_7 | 12578666 | 12580588 |
| pseudochr_OX411251.1_7 | 10490618 | 10490839 |
| pseudochr_OX411251.1_7 | 10487686 | 10488183 |
| pseudochr_OX411251.1_7 | 11401664 | 11402134 |
| pseudochr_OX411251.1_7 | 10307346 | 10307726 |
| pseudochr_OX411251.1_7 | 10499967 | 10500386 |
| pseudochr_OX411251.1_7 | 9489122 | 9491062 |
| pseudochr_OX411251.1_7 | 9876181 | 9876672 |
| pseudochr_OX411251.1_7 | 10499387 | 10499713 |
| pseudochr_OX411251.1_7 | 12331507 | 12332805 |
| pseudochr_OX411251.1_7 | 9993838 | 9994752 |
| pseudochr_OX411251.1_7 | 9167530 | 9170487 |
| pseudochr_OX411251.1_7 | 10521447 | 10524935 |
| pseudochr_OX411251.1_7 | 9862261 | 9862515 |
| pseudochr_OX411251.1_7 | 12491414 | 12492472 |
| pseudochr_OX411251.1_7 | 12512974 | 12514092 |
| pseudochr_OX411251.1_7 | 9630284 | 9632992 |
| pseudochr_OX411251.1_7 | 10497661 | 10498047 |
| pseudochr_OX411251.1_7 | 9445397 | 9445999 |
| pseudochr_OX411251.1_7 | 10501942 | 10502778 |
| pseudochr_OX411251.1_7 | 10494750 | 10495439 |
| pseudochr_OX411251.1_7 | 12072057 | 12073814 |
| pseudochr_OX411251.1_7 | 10143252 | 10146233 |
| pseudochr_OX411251.1_7 | 10501239 | 10501592 |
| pseudochr_OX411251.1_7 | 10495477 | 10495863 |
| pseudochr_OX411251.1_7 | 11009683 | 11010276 |
| pseudochr_OX411251.1_7 | 9302780 | 9304780 |
| pseudochr_OX411251.1_7 | 10492033 | 10492641 |
| pseudochr_OX411251.1_7 | 10414416 | 10415501 |
| pseudochr_OX411251.1_7 | 12455082 | 12455810 |
| pseudochr_OX411251.1_7 | 12108695 | 12109966 |
| pseudochr_OX411251.1_7 | 9319978 | 9321342 |
| pseudochr_OX411251.1_7 | 12435807 | 12437342 |
| pseudochr_OX411251.1_7 | 9324208 | 9326034 |
| pseudochr_OX411251.1_7 | 10002926 | 10004062 |
| pseudochr_OX411251.1_7 | 12467806 | 12468372 |
| pseudochr_OX411251.1_7 | 10114700 | 10115206 |
| pseudochr_OX411251.1_7 | 10525520 | 10525909 |
| pseudochr_OX411251.1_7 | 10527226 | 10527945 |
| pseudochr_OX411251.1_7 | 9967461 | 9970040 |
| pseudochr_OX411251.1_7 | 10492638 | 10493942 |
| pseudochr_OX411251.1_7 | 9251123 | 9254803 |
| pseudochr_OX411251.1_7 | 9860614 | 9862149 |
| pseudochr_OX411251.1_7 | 10514733 | 10515107 |
| pseudochr_OX411251.1_7 | 9791704 | 9793509 |
| pseudochr_OX411251.1_7 | 10496431 | 10496829 |
| pseudochr_OX411251.1_7 | 10503121 | 10504068 |
| pseudochr_OX411251.1_7 | 10119027 | 10120142 |
| pseudochr_OX411251.1_7 | 10308005 | 10308577 |
| pseudochr_OX411251.1_7 | 12425473 | 12428769 |
| pseudochr_OX411251.1_7 | 12464996 | 12465898 |
| pseudochr_OX411251.1_7 | 10495866 | 10496408 |
| pseudochr_OX411251.1_7 | 9749700 | 9750161 |
| pseudochr_OX411251.1_7 | 9277837 | 9279555 |
| pseudochr_OX411251.1_7 | 10456494 | 10456994 |
| pseudochr_OX411251.1_7 | 12456154 | 12456504 |
| pseudochr_OX411251.1_7 | 9250512 | 9251126 |
| pseudochr_OX411251.1_7 | 9255741 | 9257024 |
| pseudochr_OX411251.1_7 | 10458860 | 10460140 |
| pseudochr_OX411251.1_7 | 9308467 | 9309075 |
| pseudochr_OX411251.1_7 | 9971195 | 9972310 |
| pseudochr_OX411251.1_7 | 11430956 | 11432653 |
| pseudochr_OX411251.1_7 | 9195758 | 9196834 |
| pseudochr_OX411251.1_7 | 12548097 | 12549185 |
| pseudochr_OX411251.1_7 | 12488520 | 12489467 |
| pseudochr_OX411251.1_7 | 9496574 | 9497347 |
| pseudochr_OX411251.1_7 | 10448008 | 10449354 |
| pseudochr_OX411251.1_7 | 12465999 | 12466823 |
| pseudochr_OX411251.1_7 | 9760099 | 9761982 |
| pseudochr_OX411251.1_7 | 10528650 | 10529567 |
| pseudochr_OX411251.1_7 | 9285581 | 9287386 |
| pseudochr_OX411251.1_7 | 10499734 | 10499967 |
| pseudochr_OX411251.1_7 | 9880016 | 9882286 |
| pseudochr_OX411251.1_7 | 12446577 | 12448175 |
| pseudochr_OX411251.1_7 | 10526056 | 10526574 |
| pseudochr_OX411251.1_7 | 11402152 | 11402388 |
| pseudochr_OX411251.1_7 | 12466968 | 12467720 |
| pseudochr_OX411251.1_7 | 10517445 | 10521329 |
| pseudochr_OX411251.1_7 | 12552196 | 12553527 |
| pseudochr_OX411251.1_7 | 9479432 | 9481534 |
| pseudochr_OX411251.1_7 | 9862550 | 9862858 |
| pseudochr_OX411251.1_7 | 9747267 | 9748256 |
| pseudochr_OX411251.1_7 | 9752280 | 9753068 |
| pseudochr_OX411251.1_7 | 10439996 | 10440739 |
| pseudochr_OX411251.1_7 | 9124443 | 12716725 |
| pseudochr_OX411256.1_12 | 1750881 | 1750928 |
| pseudochr_OX411256.1_12 | 4190000 | 4190146 |
| pseudochr_OX411260.1_16 | 2505115 | 2505380 |
| pseudochr_OX411262.1_18 | 6868881 | 6869945 |
