## Supplementary figures and images for "De Novo Genome Assembly and Annotation for the Webbing Clothes Moth (*Tineola bisselliella*): A Globally Distributed, Economically Important Pest"

### Figure S1

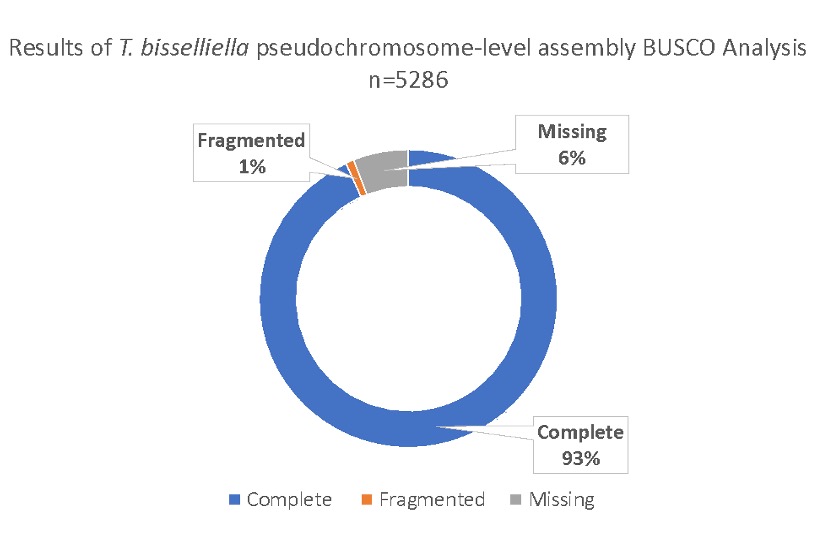

### Figure S2

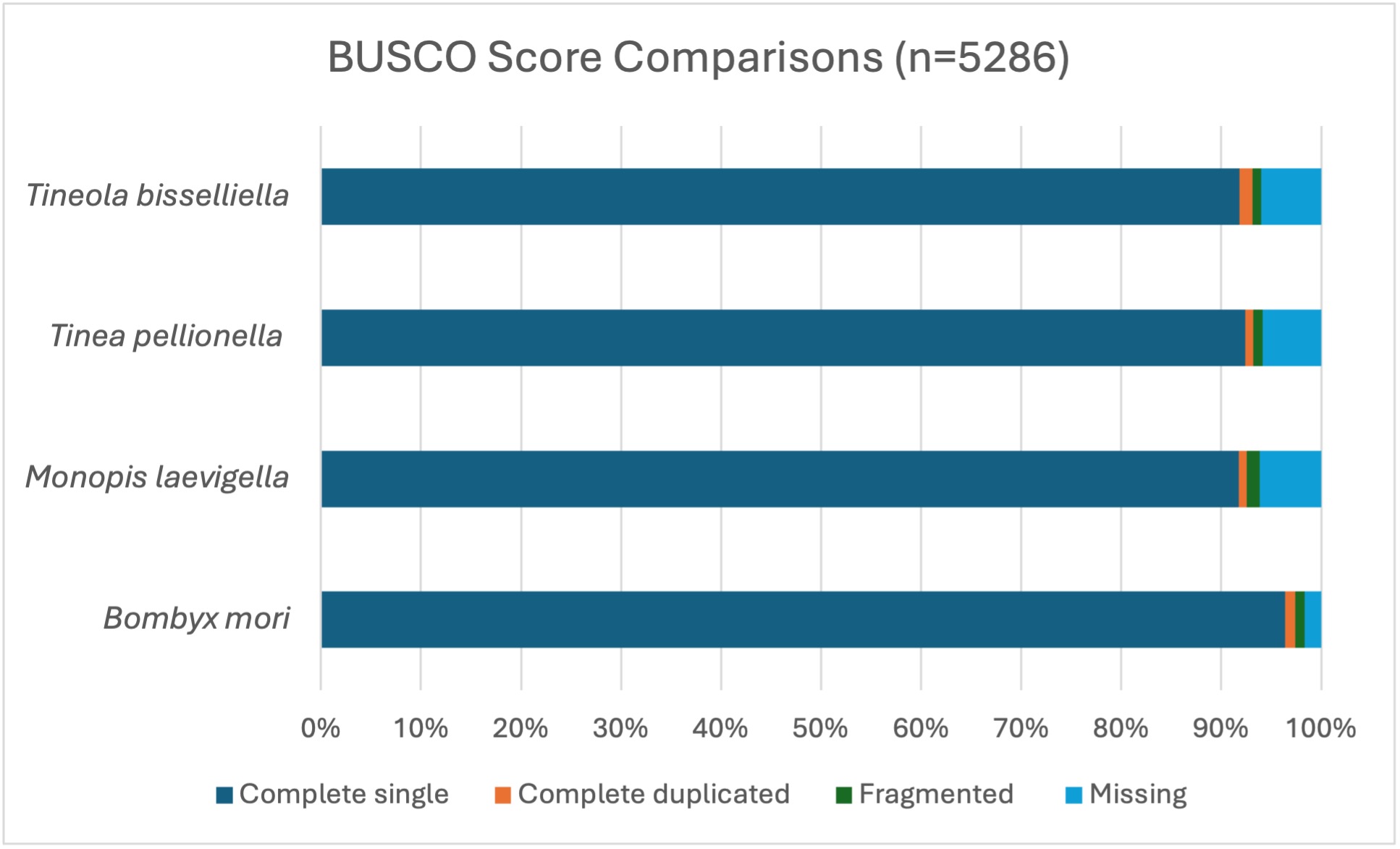

### Figure S3

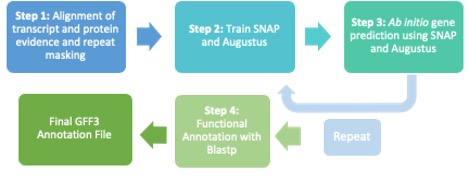

### Figure S4

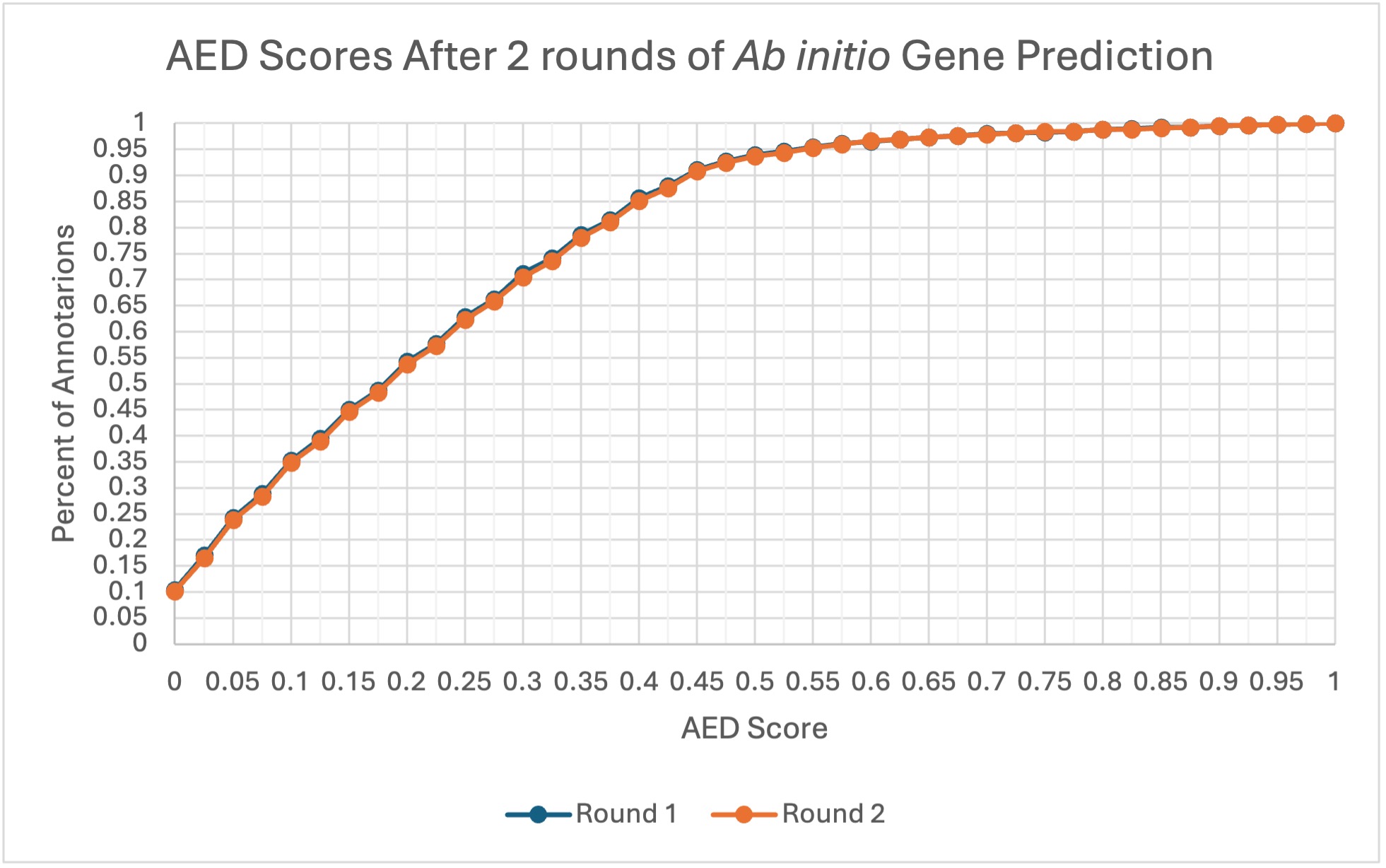

### Figure S5

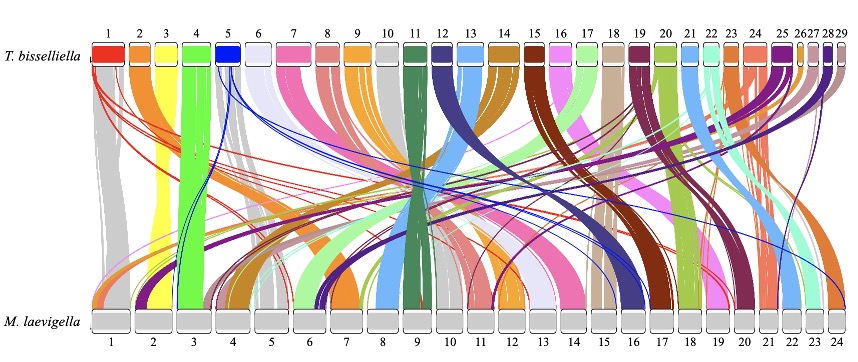

### Figure S6

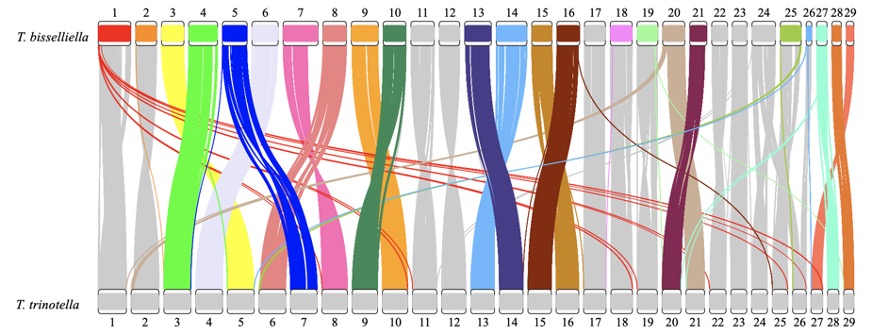

### Figure S7

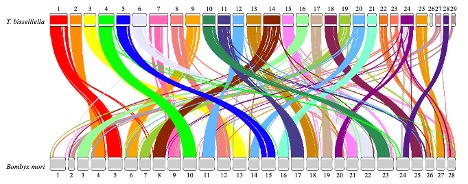
